# Phosphorylation and O-GlcNAcylation at the same α-synuclein site generate distinct fibril structures

**DOI:** 10.1101/2023.06.27.546682

**Authors:** Jinjian Hu, Wencheng Xia, Shuyi Zeng, Yeh-Jun Lim, Youqi Tao, Yunpeng Sun, Lang Zhao, Haosen Wang, Dan Li, Cong Liu, Yan-Mei Li

## Abstract

α-Synuclein (α-syn) forms amyloid fibrils that are critical in the progression of Parkinson’s disease (PD) and serves as the pathological hallmark of PD. Different posttranslational modifications (PTMs) have been identified at multiple sites of α-syn, influencing its conformation, aggregation and function. Here, we investigate how disease-related phosphorylation and O-GlcNAcylation at the same α-syn site (S87) affect fibril structure and neuropathology. Using semi-synthesis, we obtained homogenous α-syn monomer with site-specific phosphorylation (pS87) and O-GlcNAcylation (gS87) at S87, respectively. Cryo-EM analysis revealed that pS87 and gS87 α-syn form two novel but distinct fibril structures. The GlcNAc situated at S87 establishes interactions with K80 and E61, inducing a unique iron-like fold with the GlcNAc molecule on the iron handle. While, phosphorylation at the same site prevents a lengthy C-terminal region including residues 73-140 from incorporating into the fibril core due to electrostatic repulsion. Instead, the N-terminal half (1-72) shapes a novel arch-like fibril structure. We further show that both pS87 and gS87 α-syn fibrils display reduced neurotoxicity and propagation activity compared with unmodified α-syn fibril. Our findings demonstrate that different PTMs at the same site can produce distinct fibril structures, which emphasizes the precise regulation of PTMs to amyloid fibril formation and pathology.

## INTRODUCTION

Pathological aggregation of amyloid proteins such as α-syn and Tau is strongly linked to neurodegenerative diseases (NDs)^1-3^. Numerous posttranslational modifications (PTMs), such as phosphorylation, O-glycosylation, ubiquitination, and acetylation, have been identified on amyloid proteins, playing diverse roles in regulating protein conformation, aggregation kinetics, fibril structures and pathology^4-9^. α-Syn is the key player in Parkinson’s disease (PD), with its amyloid fibril being the main component of the Lewy bodies (LBs) and Lewy neurites (LNs), hallmarks of PD^10-12^. Elevated α-syn phosphorylation level, including pY39, pS87, and pS129, has been observed in the brains of PD and MSA patients^13-17^. While pY39 and pS129 are dominant PTMs in LBs and LNs, pS87 is not enriched^13, 15, 18^. Moreover, unlike pY39 and pS129 which enhances α-syn transmission and pathology in cells^7, 19^, pS87 has been found to prevent α-syn aggregation and neurotoxicity^14, 20^. This suggests that phosphorylation at different α-syn sites may have distinct or even opposing roles in regulating α-syn aggregation and pathology.

In addition to phosphorylation, amyloid proteins such as Tau and α-syn are found to be O-GlcNAcylated^21-23^, which may prevent their amyloid aggregation^8, 24^. Interestingly, there is a noticeable reduction in the overall O-GlcNAc levels in brains affected by Alzheimer’s Disease^25-26^. Neurodegeneration in mice is also observed when the overall O-GlcNAc level is diminished through the suppression of O-GlcNAc transferase^27^. This suggests that O-GlcNAcylation could have a protective function in NDs. It’s worth mentioning that α-syn was observed to be O-GlcNAcylated at several points including T72, T75, T81, and S87^22, 28-29^, and this consistently demonstrated protection against α-syn aggregation^8, 24, 30-32^. Notably, T75 and S87 are sites of both phosphorylation and O-GlcNAcylation^14, 28-29^. However, the specific effects of different PTMs at the same α-synuclein sites in influencing amyloid aggregation and neuropathology are not yet fully understood.

In this study, we focus on α-syn S87, which can be modified by both phosphorylation and O-GlcNAcylation. Employing a semi-synthesis approach, we prepared pS87 and gS87 α-syn monomer with high purity. Remarkably, we revealed by using cryo-EM that pS87 α-syn and gS87 α-syn form amyloid fibrils with distinct structures. In gS87 α-syn fibril structure, GlcNAc at S87 establishes new interactions with neighboring residues in the C-terminal of the non-amyloid-β component (NAC) of α-syn, inducing a novel iron-like structure. While, in pS87 α-syn fibril structure, phosphate group excludes the C-terminal of α-syn NAC from forming fibril core. Moreover, both pS87 α-syn and gS87 α-syn fibrils exhibit reduced neuropathology compared to the unmodified Wild-type (WT) α-syn fibril formed under the same condition. Our findings provide structural basis of how different PTMs lead to the formation of distinct fibril structures with attenuated neurotoxicity.

## RESULTS

### Semi-synthesis of pS87 and gS87 α-syn

We first sought to obtain site-specific pS87 and gS87 α-syn with high purity using native chemical ligation strategy using peptide hydrazides developed by Liu and co-workers^33-34^. Since S87 is in the middle of α-syn, we performed N-to-C sequential native chemical ligation using three segments referring to the work of Pratt and co-workers^8, 31, 35^, including segment X (α-syn 1-84 thioester), segment Y (gS87/pS87 α-syn A85C-90NHNH_2_), segment Z (α-syn A91C-140) (Figure 1a).

**Figure 1.**
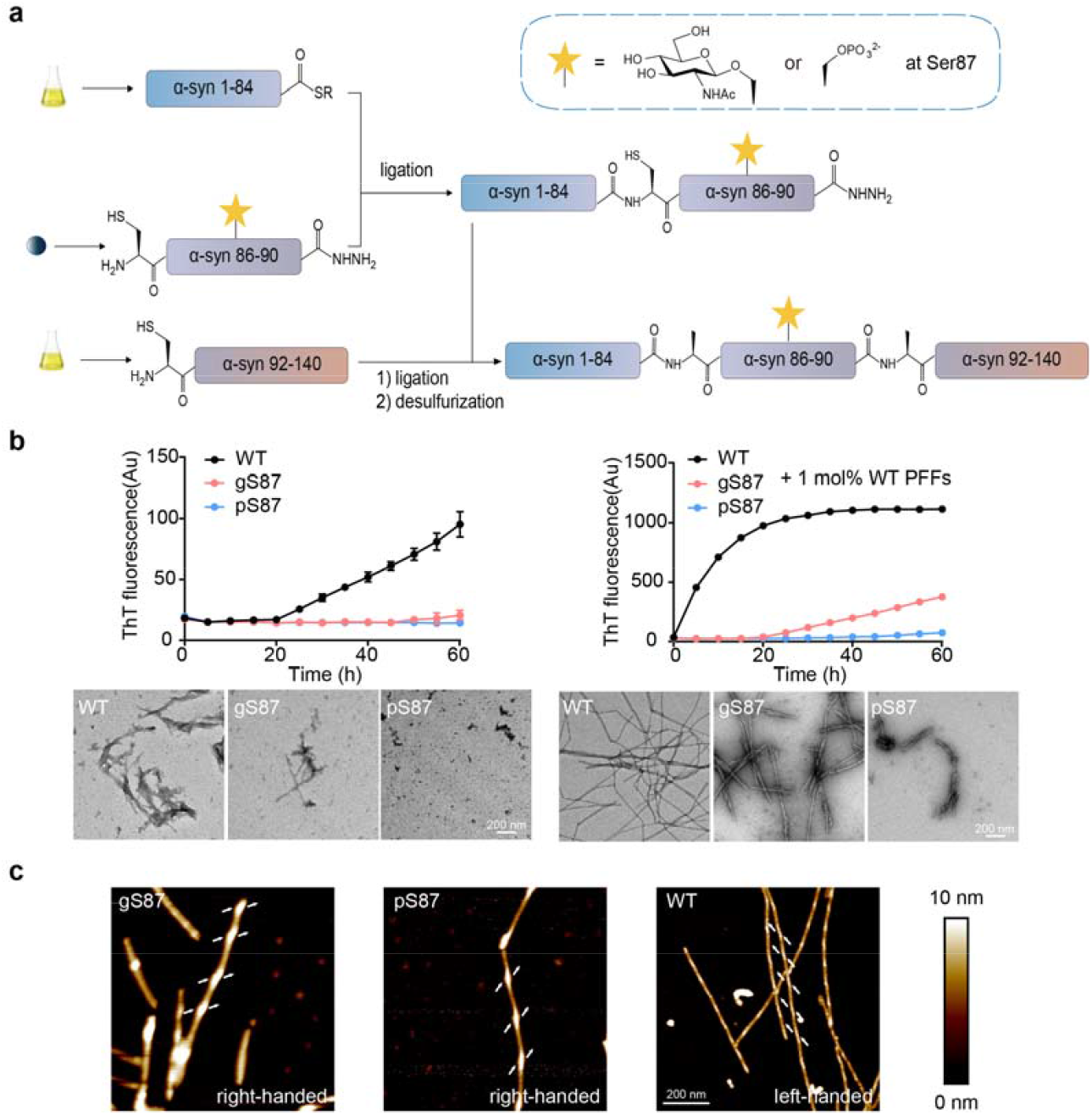
Synthesis workflow and fibril characterization of gS87, pS87 and unmodified WT α-syn. (a) Workflow of the semi-synthesis of α-syn with different modifications at S87. (b) Left: ThT kinetic assay (top) and NS-TEM images (bottom) of unmodified WT, gS87, and pS87 α-syn fibrils. Right: ThT kinetic assay (top) and NS-TEM images (bottom) of unmodified WT, gS87 and pS87 α-syn fibrils in the presence of 1 mol% PFF formed by the unmodified WT α-syn monomer. The fibrils were characterized by NS-TEM at the endpoint (60h) of the ThT kinetic assay. Data correspond to mean ± s.d., n=3. Scale bar: 200 nm. (c) AFM images of gS87 α-syn, pS87 α-syn, and unmodified WT α-syn fibril. The arrows at both sides of the fibril indicate the starting points of the fibril protrusions to clarify the handedness. Scale bar: 200 nm.

To obtain segment X, α-syn 1-84 fused with intein was expressed, then followed by thiolysis. Segment α-syn Y-gS87 and α-syn Y-pS87 were manually synthesized by Fmoc-based solid phase peptide synthesis. The glycosylated Fmoc-amino acid, Fmoc-L-Ser(GlcNAc(Ac)_3_-β-D)-OH was synthesized referring to the work of Pratt^36^. The recombinant segment Z was expressed in E. *coli*. Methylhydroxylamine was then used to reverse N-terminal cysteine modification during expression.^37^ Segment Y and Z were further purified and lyophilized for next reaction.

After obtaining these segments, we firstly ligated X and Y using native chemical ligation^38^, followed by purifying. The resulting segment XY-gS87/pS87 (gS87/pS87 α-syn 1-90NHNH_2_, A85C) was lyophilized and ligated with segment Z through protein hydrazide. After purification and lyophilization, XYZ with gS87/pS87 modification (gS87/pS87 α-syn, A85C A91C) was obtained. To remove sulfhydryl groups on the cysteine residues at the ligation sites, radical-catalyzed desulfurization^39^ was performed to obtain α-syn protein only with modification at S87. All segments during synthesis and the final modified protein products were characterized with analytical RP-HPLC and ESI-MS (Figure S2-4).

### Characterization of the gS87 and pS87 α-syn fibrils

We next investigated the influence of glycosylation and phosphorylation at S87 on α-syn fibrillation through ThT kinetic assay and negative-staining (NS) transmission electron microscopy (TEM). As shown in Figure 1b, both gS87 and pS87 α-syn exhibited significantly reduced capability for fibrillation compared to unmodified WT α-syn. In the presence of pre-formed fibrils (PFFs) formed by unmodified α-syn protein, unmodified monomer rapidly formed fibril without nucleation (Figure 1b). In sharp contrast, gS87 α-syn started to form fibril after 20 hours incubation. The ThT signal of pS87 sample slowly picked up after 40 hours with much less fibril formed as revealed by NS-TEM (Figure 1b). More interestingly, atomic force microscopy (AFM) revealed that both the gS87 α-syn and pS87 α-syn fibrils feature a right-handed helical twist, which is distinct from the left-handed unmodified WT α-syn fibril (Figure 1c). Together, our results demonstrate that both glycosylation and phosphorylation at S87 severely impair α-syn fibrillation, and induce α-syn to form distinct morphological fibrils from unmodified fibril.

### Cryo-EM structure determination of gS87 and pS87 α-syn fibrils

To investigate whether glycosylation and phosphorylation at S87 alters the structures of α-syn amyloid fibrils, we set out to determine the atomic structure of gS87 and pS87 α-syn fibril by using cryo-EM. The cryo-EM data were collected on a Titan Krios G4 cryo-transmission electron microscope (2,134 micrographs for gS87 and 2,423 micrographs for pS87). 21,328 fibrils from gS87 dataset and 27,806 fibrils from pS87 dataset were picked for the following two-dimensional (2D) classification and 3D reconstruction (Table S1).

For gS87 sample, 2D classification results reveal there are two fibril polymorphs in the gS87 α-syn fibril, including the double filament polymorph (∼56%) and the single filament polymorph (∼44%) (Figure S5a). After performing the 3D reconstruction, we managed to obtain the density map of the double filament with the overall resolution of density map of 3.1 Å (Figure S5c). The double filament of gS87 fibril features a half pitch (a whole pitch represents the length of a 360° helical turn of the entire fibril) of ∼156 nm, a helical rise of 2.41 Å, and a helical twist of -179.72° (Figure 2a). gS87 α-syn adopts a similar architecture in both the double filament and single filament polymorphs (Figure S5a). For pS87 sample, 2D classification identified two different fibril morphologies, including the straight filament polymorph (∼74%) and the twisted filament polymorph (∼26%) (Figure S5b). We managed to obtain the high-quality density map of the twisted filament with an overall resolution of 2.6 Å (Figure S5d). The pS87 twisted filament features a half pitch of ∼154 nm, a helical rise of 2.41 Å, and a helical twist of -179.72° (Figure 2b).

**Figure 2.**
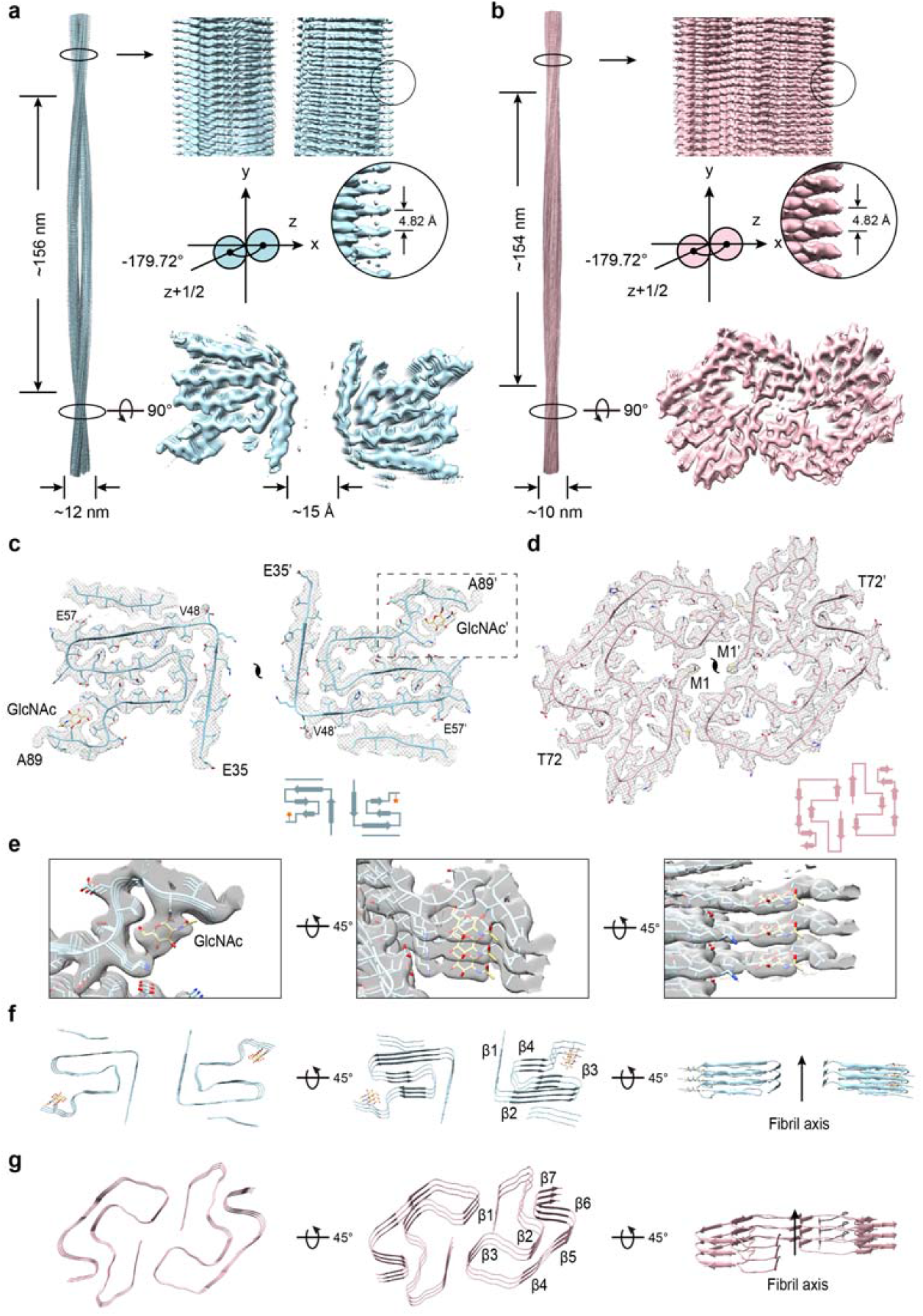
Cryo-EM structures of gS87 and pS87 fibrils. (a-b) The density maps of the gS87 (a) and pS87 (b) fibril are colored in light-blue and pink, respectively. Fibril parameters including half-pitch, fibril width, twist angle, and helical rise are marked. (c-d) Cross-section view for the density maps with a built-in structure model of gS87 (c) and pS87 (d) α-syn. Topology diagrams are shown at the bottom right. (e) Zoom-in views of the GlcNAc molecules in the electron density from (c) are shown. (f-g) Views of three layers of gS87 (f) and pS87 (g) α-syn fibrils are shown in the cartoon. The β-strands of the fibril structures are numbered and labeled accordingly with the fibril axis indicated.

### Structural analysis of gS87 and pS87 α-syn fibrils

Utilizing high-quality cryo-EM density maps, we constructed two structural models for the double filament polymorph of gS87 (hereafter referred to as gS87 α-syn fibril) and the twisted filament polymorph of pS87 (hereafter referred to as pS87 α-syn fibril). Remarkably, α-syn exhibits two entirely distinct and previously unobserved structural conformations in these two fibril structures (Figure 2c, d). In the gS87 α-syn fibril, α-syn adopts a conserved iron-like fold in the core structure with the modified GlcNAc molecule on the iron handle (Figure 2c). The superior electron density of GlcNAc suggests its high stability within the core structure (Figure 2e). The GlcNAc molecules stack on top of each other and align along the fibril axis with an intermolecular distance of 4.82 Å. The fibril core comprises residues E35 to A89, which fold into 4 β-strands (β1-β4) (Figure 2f). An unassigned island is observed on the outer surface of the fibril core, adjacent to β2 (residues 48-57) (Figure S6a). The two identical protofilaments intertwine to form the gS87 fibril (Figure 2c). Intriguingly, unlike the tight and complementary protofilament interface seen in earlier fibril structures^7, 40-43^, neighboring α-syn chains from the two protofilament sets remain separate, without any direct interaction. The distance between the nearest side chains of the two α-syn molecules exceeds ∼15 Å (Figure 2a), suggesting a relatively weak interaction between the two protofilaments.

In the pS87 α-syn fibril, α-syn adopts an arch-like fold in the fibril core, composed of residues M1 to T72 folding into 7 β-strands (β1-β7) (Figure 2g). Unlike the gS87 and unmodified WT α-syn fibrils formed under identical condition^40^, the entire N-terminal region participates in fibril core formation in the pS87 fibril (Figure 2d). While, the phosphorylated S87 is excluded from the fibril core, and remains flexible and invisible from cryo-EM. Furthermore, the two protofilaments of pS87 α-syn are zipped together through extensive intermolecular interactions involving the side chains of residues V3, M5, L38 and V40 (Figure 2d). In summary, although phosphorylation and O-glycosylation modify the same residue, they lead to substantially distinct conformation of the α-syn subunit and protofilament arrangement in the fibril structures.

### Comparison of α-syn fibril structures with two different modifications

In the gS87 α-syn fibrils, the GlcNAc installed at S87 participates in fibril core formation and is located at the C-terminus of the core structure (Figure 2c). GlcNAc forms direct interactions with the K80, T81, V82, I88 and A89 (Figure 3a). Concurrently, K80 establishes a salt bridge with E61 (Figure 3a), stabilizing the U-shaped structure formed by β3 and β4. Additionally, hydrophilic zipper-like interactions formed by residues 61-65 and 79-80 contribute to the stabilization of the U-shaped structure (Figure S6b). β3 connects to the two U-shaped structures through hydrophobic zipper-like interactions with β2, mediated by residues 66-71 (Figure 3a). β1 further wraps around the U-shaped structures, forming the iron-like core of S87 (Figure S6c). Intriguingly, despite that unmodified WT α-syn employs the same region (residues 37-99) for forming fibril core^40^, it features varying numbers of β-strand formed by different segments, which assemble into a unique Greek key-like structure (Figure 3b, 4a, b). In the unmodified structure, S87 does not form any direct interactions with other residues (Figure 3b). While, K80, which forms an interaction with GlcNAc at S87 in the gS87 fibril structure, establishes a salt bridge with E46 at the edge of the Greek key (Figure 3b). This K80-E46 salt bridge is essential for maintaining the entire Greek key-like structure, and its disruption by the E46K mutation effectively abolishes the Greek key-like fold^41^. Therefore, the GlcNAc modified at S87 introduces new interactions with K80 and E61, leading to structural rearrangement of α-syn and the formation of a distinct β-strand pattern to create a new fibril core structure (Figure 4a, b).

**Figure 3.**
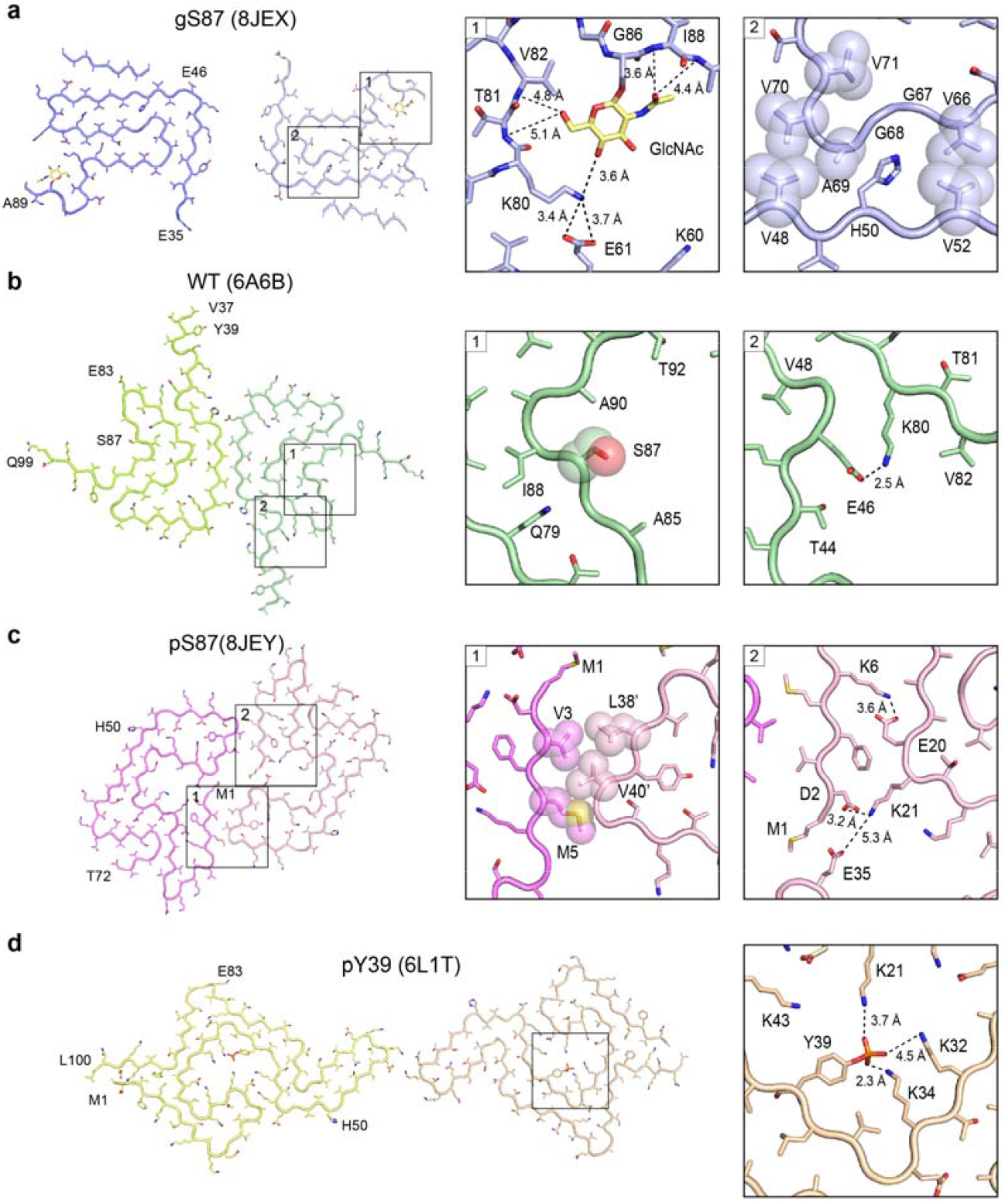
Structural analysis of gS87, pS87, unmodified WT, and pY39 α-syn fibril. (a) The structural model of gS87 fibril, with the zoom-in views the interactions between GlcNAc, K80, E61 T81, V82, I88 and A89, and the hydrophobic zipper-like interactions with the involved residues labeled. (b) The structure of unmodified WT α-syn fibril, with the conformation of S87 and the salt bridge between K80 and E46 shown in the zoom-in views. (c) The structure of pS87 fibril, with the hydrophobic interactions of the interface between two protofilaments, and two pairs of salt bridges shown in the zoom-in views. Residues involved in the inter-protofilamental interactions are shown in spheres. (d) The structure of pY39 fibril with the electrostatic interactions of the phosphate group bound to Y39 and K21, K32 and K34 shown in the zoom-in view. Distances are shown in Å. The PDB code of each fibril structure is indicated.

**Figure 4.**
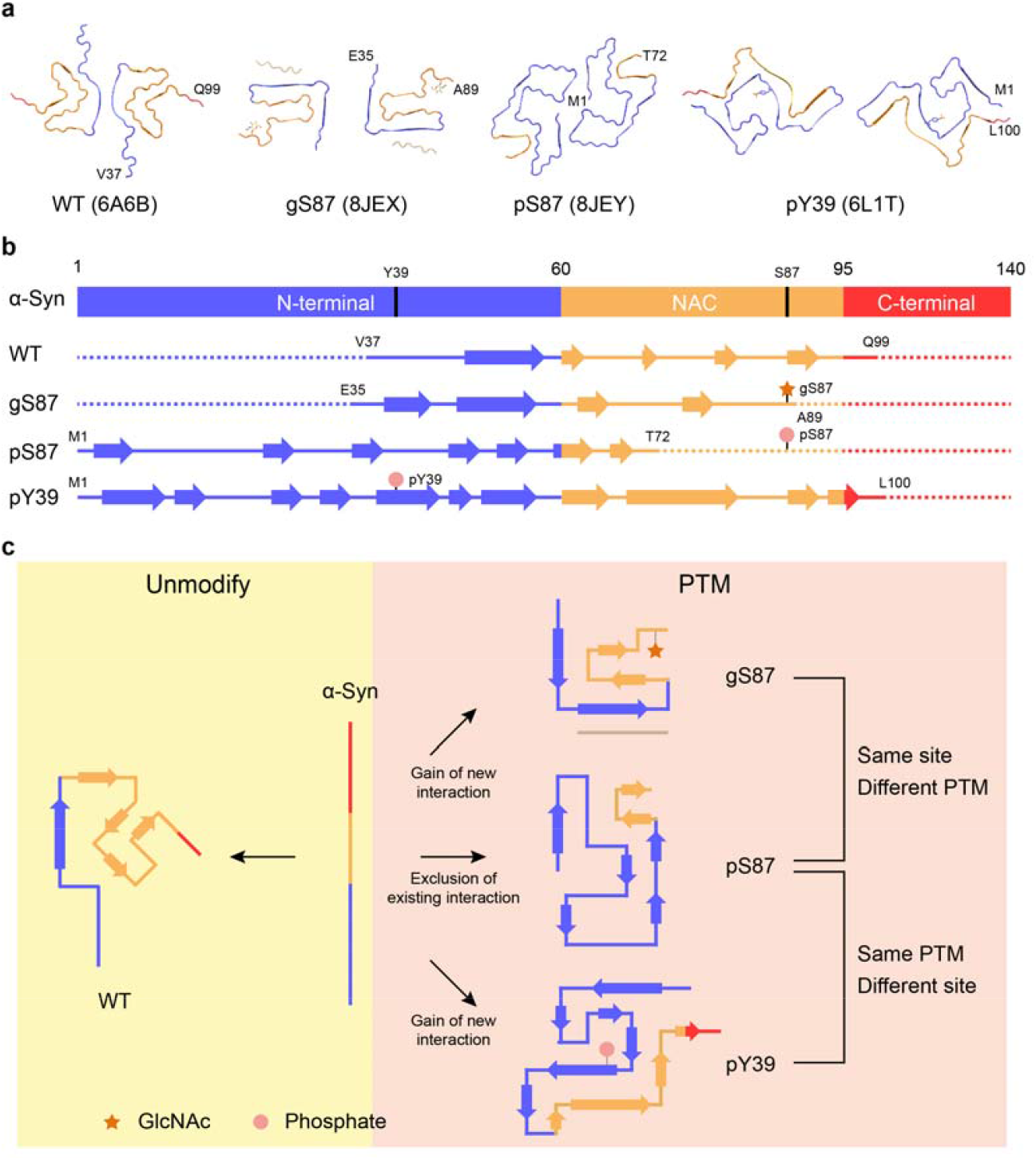
Comparison of α-syn fibril structures with different modifications. (a) one-layer models of unmodified WT (PDB: 6A6B), gS87 (PDB:8JEX), pS87 (PDB:8JEY) and pY39 (PDB: 6L1T) fibrils with the N-terminal region colored in blue, the NAC colored in yellow and the C-terminal region colored in red. (b) The secondary structure alignment of four α-syn fibril structures from (a) with different colors for three regions. (c) Schematic diagram shows that two different PTMs modified at different sites induce three distinct fibril core structures.

Remarkably, when phosphorylated, S87 is excluded from the fibril core, contrasting sharply with its O-glycosylated counterpart in the fibril structure (Figure 3a). Starting from residue 73, the extended C-terminal region (residues 73-140) remains highly flexible in the fibril structure, without electron density. Conversely, the entire N-terminal region (residues 1-35), absent in both gS87 and unmodified α-syn fibril cores, is incorporated into the pS87 fibril core (Figure 3c). In the pS87 fibril structure, residues 1-29 from the N-terminal form a U-shaped structure stabilized by two pairs of salt bridges, including K6 & E20 and K21 & D2 (Figure 3c). Meanwhile, V3 and M5 on β1 establish hydrophobic interactions with residues L38’ and V40’ from the neighboring chain, forming intermolecular interactions that bring the two protofilaments together (Figure 3c). The second U-shaped structure is formed by residues 30-49, stabilized by a salt bridge between K32 and E46 and a hydrogen bond between K34 and Y39 (Figure S6d). β2 (residues 26-29) forms a steric zipper-like hydrophobic interaction with residues 52-55, connecting the two U-shaped structures (Figure S6d).

Intriguingly, our prior structural results demonstrated that phosphorylation at Y39 of α-syn enables the integration of the entire N-terminal region into the fibril core^7^ (Figure 3d). In the pY39 fibril structure, the phosphate group modified at Y39 establishes extensive electrostatic interactions with K21, K32, and K34 (Figure 3d). These interactions are vital for incorporating the entire N-terminal region into the fibril core, resulting in a large hook-like architecture (Figure 4a, b). This structure suggests that if a phosphate group participates in fibril core formation, it needs to be stabilized by positively charged residues^7^. However, in the pS87 fibril structure, the positively charged residues are mostly localized in the N-terminal region, which either form electrostatic interactions with negatively charged residues (K6 & E20, K21 & D2, K32 & E46) or are situated on the fibril’s outer surface (Figure 3c). Consequently, the addition of a phosphate group to S87 prevents the C-terminal region of NAC from participating in fibril core formation due to electrostatic repulsion, obstructing α-syn’s ability to form the Greek key-like fold observed in unmodified WT α-syn fibril. In contrast, the entire N-terminal region is incorporated into the fibril core, giving rise to a unique arch-like architecture in the pS87 fibril.

### gS87 and pS87 α-syn fibrils exhibit reduced neurotoxicity and propagation activity

Finally, we investigated whether the novel fibril structures induced by glycosylation and phosphorylation at S87 display altered neuropathological activities using a well-established neuronal propagation model^44-46^ (Figure 5a). We added unmodified WT, gS87, and pS87 α-syn PFFs into the culture medium of rat primary cortical neurons to seed the aggregation of endogenous α-syn. After 14 days of PFFs treatment, we detected the formation of the PFF-induced endogenous α-syn aggregate with an antibody of pathological pS129 α-syn through immunofluorescence staining. Remarkably, both groups treated with gS87 and pS87 α-syn PFFs displayed a significant reduction in pathological pS129 α-syn propagation compared to the unmodified WT α-syn PFFs-treated group (Figure 5b, c). The pS87 PFFs exhibited the lowest propagation activity (Figure 5c). Moreover, we measured the cytotoxicity of unmodified WT, gS87, and pS87 α-syn fibrils to neurons using a cell counting kit-8 (CCK-8) assay. The result showed that both gS87 and pS87 α-syn PFFs exhibited considerably decreased toxicity to primary neurons compared to unmodified WT α-syn PFFs, with pS87 PFFs displaying the lowest neurotoxicity (Figure 5d). In conclusion, our findings indicate that glycosylation and phosphorylation at S87 induce α-syn to form two distinct fibril structures with reduced propagation activity and neurotoxicity in neurons.

**Figure 5.**
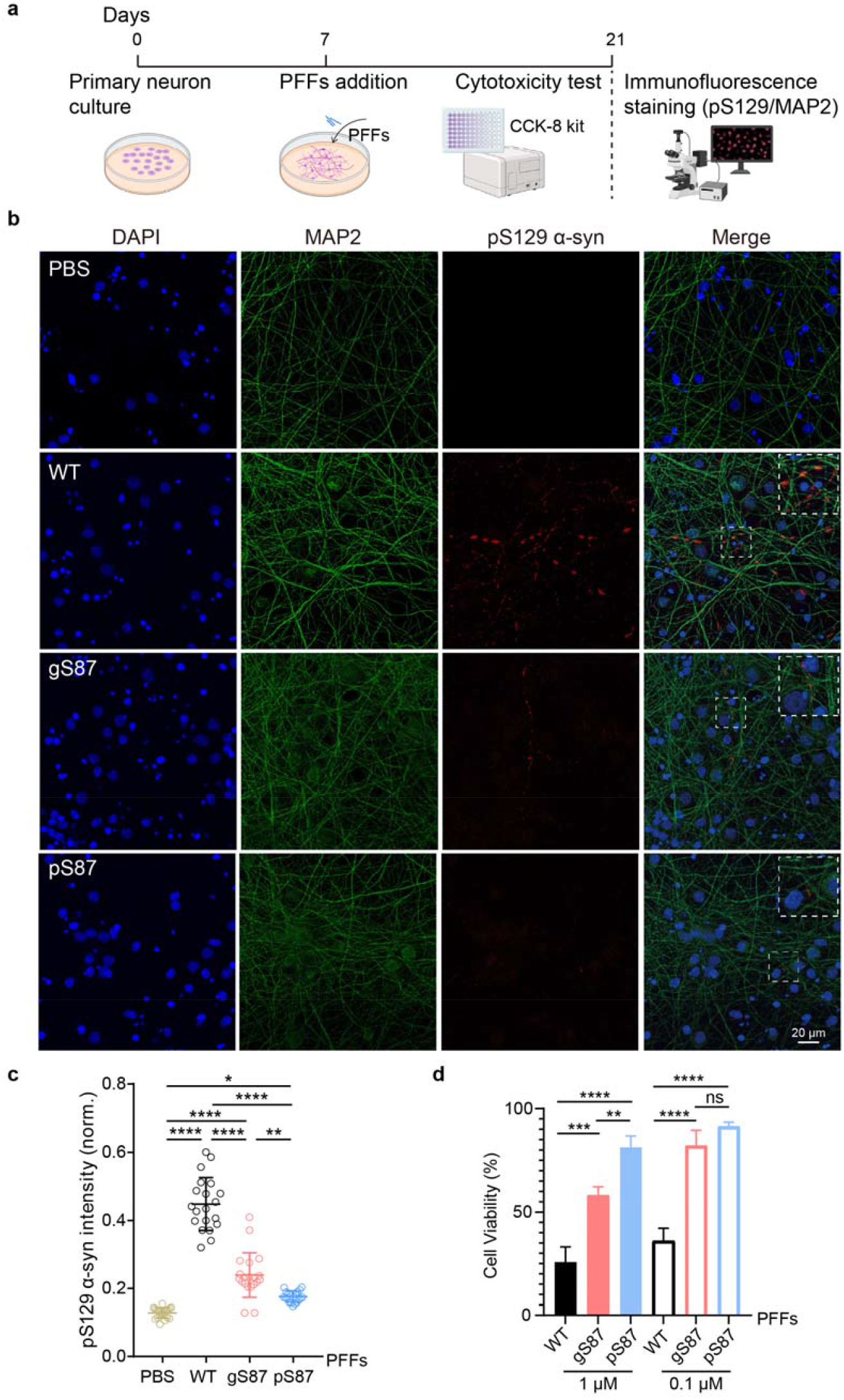
Neurotoxicity and propagation activity measurement of the unmodified WT, gS87 and pS87 α-syn fibrils. (a) Workflow of cytotoxicity test and immunofluorescence staining in rat primary neurons. (b) Representative immunofluorescence images of rat primary neurons treated with PBS, 100 nM unmodified WT PFFs, gS87 PFFs and pS87 PFFs for 14 d, respectively. DAPI (blue), pS129 α-syn (red) and microtubule-associated protein 2 (MAP2) (green) were stained. Scale bar: 20 μm. Images were processed by Image J. (c) Quantitative analyses of pS129 α-syn aggregation induced by different types of fibrils. The intensity of pS129 α-syn was normalized to the intensity of DAPI for each sample. One-way ANOVA followed by Tukey’s post-hoc test. Data shown are mean ± s.d.. * p<0.05, ** p<0.01, **** p<0.0001. (d) Cytotoxicity measurement of the unmodified WT, gS87 and pS87 PFFs using CCK-8 kit. One-way ANOVA followed by Tukey’s post-hoc test. Data shown are mean ± s.d., n = 3 independent samples (biological repetitions). ns, p>0.05, ** p<0.01, *** p<0.001, **** p<0.0001.

## DISCUSSION

Various types of PTMs have been identified that modify α-syn at different sites, playing diverse roles in regulating α-syn monomer conformation, membrane binding, amyloid aggregation kinetics and the atomic structures and pathology of α-syn fibril^7, 28^. In this study, we explore whether distinct PTMs at the same residue can lead to the formation of different fibril structures. Intriguingly, we found that O-GlcNAcylation at the S87 residue can introduce new intra-molecular interactions, resulting in the formation of a novel iron-like fold in the fibril structure (Figure 3a, 4a). This O-GlcNAcylation generates a new fibril polymorph by directly establishing new interactions. In contrast, phosphorylation at the same site prevents the involvement of the C-terminal region of NAC in the fibril core due to electrostatic repulsion. Instead, it promotes the incorporation of the entire N-terminal region to form a previously unidentified, enlarged arch-like fibril core structure (Figure 3c, 4a). Therefore, this phosphorylation generates a new fibril polymorph by excluding certain NAC region from forming fibril core. Consequently, different PTMs at the same site may utilize entirely distinct mechanisms to direct fibril core formation, depending on the specific properties of the modified groups (Figure 4c).

Additionally, we compared the fibril structures of pS87 α-syn with pY39 α-syn, and found that the same phosphate group installation at different sites can also produce completely different fibril structures (Figure 3c, d, 4a). In pY39 α-syn, the phosphate group added at Y39 forms extensive interactions with three lysine residues in the N-terminal region, inducing the formation of a large hook-like fibril structure (Figure 3d, 4a). However, adding the phosphate group at S87 results in the exclusion of the C-terminal region of NAC from incorporation into the fibril core due to electrostatic repulsion. Thus, the local context of the modified site may directly impact the regulation mechanism of a given modified group, adding an additional layer of complexity to the regulation of fibril structural polymorphs by PTMs (Figure 4c).

Notably, both gS87 and pS87 α-syn fibrils display considerably lower neurotoxicity and propagation activity compared to the unmodified WT α-syn fibril (Figure 5b-d). This finding aligns with previous observations that both pS87 and gS87 modifications protect against α-syn aggregation and pathology in cells^8, 14^. In stark contrast, the pY39 modification exacerbates α-syn pathology in cells and produces a fibril polymorph with heightened neurotoxicity^7^. However, the direct relationship between the structural polymorphs and their varying neuropathology is not yet well-established. Further investigation is necessary to understand how gS87 and pS87 fibril structures lead to reduced neuropathology. Furthermore, since both PTMs share the same modification site on α-syn, exploring the cross-talk and potential competition between these two modifications at S87 during various physiological or pathological processes, and its implication in disease progression, is of importance.

## CONCLUSIONS

PTMs affect α-syn aggregation and the associated neuropathology. Even though PTMs like phosphorylation and O-GlcNAcylation are known to occur on α-syn, the unique effects of different PTMs at identical sites remain incompletely understood. In this study, we generated pS87 and gS87 α-syn, specific to individual sites, by merging chemical synthesis with bacterial expression. The Cryo-EM structural analysis revealed that different PTMs at the same α-syn site can lead to the formation of entirely distinct fibril structures through different mechanisms. Additionally, both gS87 and pS87 α-syn fibrils demonstrated reduced neuropathology in rat cortical neurons compared to the unmodified WT α-syn fibril. These findings advance our molecular understanding of how diverse PTMs at the same site can precisely regulate the formation of amyloid fibrils.

## Data Availability

Density maps of double filament of gS87 α-syn fibril and twisted filament pS87 α-syn fibril are available in Electron Microscopy Data Bank (EMDB) with entry codes: EMD-36202 for double filament of gS87 α-syn fibril and EMD-36203 for twisted filament pS87 α-syn fibril. And the structure models have been deposited in the Protein Data Bank (PDB) with entry codes: 8JEX for double filament of gS87 α-syn fibril and 8JEY for twisted filament pS87 α-syn fibril.

## Supporting Information

Methods, materials and additional results (analytical HPLC result, ESI-MS characterization, fibril structure parameters and models analysis).

## Acknowledgements

We thank the Cryo-EM Microscopy center at Interdisciplinary Research Center on Biology and Chemistry, Shanghai Institute of Organic Chemistry for help with data collection. This work was supported by the National Natural Science Foundation of China (Grant No. 92053108, 82188101, 32170683, and 32171236,), the National Key R & D Program of China (2018YFA0507600, 2019YFA0904200, 2019YFE0120600), the Science and Technology Commission of Shanghai Municipality (STCSM) (Grant No. 20XD1425000, 2019SHZDZX02 and 22JC1410400), the Shanghai Pilot Program for Basic Research – Chinese Academy of Science, Shanghai Branch (Grant No. CYJ-SHFY-2022-005).

## Author Contributions

^†^J.H., W.X., S.Z., and Y.-J.L. contributed equally to this work.

