## Supplementary material for "Phosphorylation and O-GlcNAcylation at the same α-synuclein site generate distinct fibril structures": Figure S

**METHODS AND MATERIALS**

**Semi-synthesis of gS87 and pS87 α-syn**

The native chemical ligation strategy using peptide hydrazides of Liu and co-workers^1-2^ was used to synthesize α-syn with PTMs. We performed N-to-C sequential native chemical ligation using three segments referring to the work of Pratt and co-workers^3-5^, including segment X (α-syn 1-84 thioester), segment Y (pS87/gS87 α-syn A85C-90NHNH_2_), segment Z (α-syn A91C-140) (Figure 1a).

To obtain segment X, α-syn 1-84 was introduced into pTWIN1 vector (NEB) containing intein and chitin binding domain. BL21(DE3) *E. coli* was transformed and then cultured in Luria-Bertani (LB) broth with 50 μg/mL ampicillin sodium salt. Protein expression was induced by addition of 1 mM isopropyl-β-D-1-thiogalactopyranoside (IPTG) and incubation of cells for 20 h at 16 °C. Then, the cells were harvested in buffer (50 mM Hepes, 500 mM NaCl, pH 7.5) and lysed by ultrasonication (twice on ice, each for 20 min). The lysate was centrifuged (14 000 rpm, 90 min), and the supernatant was loaded on chitin column (NEB). After cleavage in 250 mM mercaptoethanesulfonate (MesNa), the protein thioester was concentrated by ultrafiltration (Figure S2a).

Segment α-syn Y-pS87 (CG[pS)IAA-NHNH_2_) and Segment α-syn Y-gS87 (CG[gS)IAA-NHNH_2_) were manually synthesized using 2-Cl-(Trt)-NHNH_2_ resin as described^1^. Fmoc-L-Ser (GlcNAc(Ac)_3_-β-D)-OH was synthesized referring to the work of Pratt^6^ and characterized with RP-HPLC, ESI-MS, and NMR (Figure S1). Fmoc-L-Ser(GlcNAc(Ac)_3_-β-D)-OH was coupled on peptide using 1.8eq HATU, 2eq HOAT, 5eq N-methylmorpholine (NMM) in N-Methylpyrrolidone (NMP). And the deprotection of O-acetyl groups on GlcNAc was carried out using 60% hydrazine hydrate in CH_3_OH (v/v) on resin for 15 min. Reagent K was used in cleavage. The peptides were purified and characterized with RP-HPLC (YMC-Pack ODS-A column) and ESI-MS (Figure S2b, c).

For segment Z, BL21(DE3) *E. coli* was transformed with pET-22b vector containing α-syn A91C-140. After incubation and expression. The protein was purified using osmotic-shock strategy.^7^ Briefly, after centrifuging, the collected cells were treated with osmotic shock buffer (30 mM Tris-HCl, 40% sucrose (w/v), and 2 mM EDTA, pH 7.4). After centrifugation (12 000 rpm, 20 min), cells were suspended in cold water with saturated MgCl_2_ added. Then the supernatant was collected. Extra precipitation (with HCl to pH 3.5 and NaOH to pH 7 in sequence) was also applied to increase purity. 400 mM O-methylhydroxylamine and 20 mM Tris(2-carboxyenthyl)phosphine (TCEP) was then added and incubated for 5h to reverse N-terminal cysteine modification.^8^ α-syn A91C-140 was further purified using RP-HPLC with proteonavi column (OSAKA SODA), then lyophilized for next reaction.

Then we ligated X and Y. Briefly, X thioester (2.5 mM, 1 eq.) and Y-pS87/Y-gS87 (2 eq.) were dissolved in ligation buffer (6 M guanidine-HCl, 200mM phosphate buffer, pH 7.0). Then, 50 eq. TCEP and 4-mercaptophenylacetic acid (MPAA) (in 200 mM phosphate buffer, pH 6.8) were added. The solution was shacked at 30 °C for 5h, then purified using RP-HPLC with proteonavi column. The product XY-pS87/gS87 was lyophilized for next reaction (Figure S2d).

For the ligation of XY and Z, segment XY (5 mM, 1eq.) was dissolved in 6 M guanidine-HCl, 200 mM phosphate buffer at pH 3.0 and put into a -15~-20 °C ice–salt bath. 0.5 M NaNO_2_ (10 eq.) was added in the reaction or 15 min. After that, 40 eq. MPAA and 1.2 eq. segment Z in buffer (200 mM phosphate buffer, 6 M guanidine-HCl, pH 6.5) was added and then the pH was adjusted to 6.8. The final concentration of XY is about 2.5 mM. The mixture was shaken at 16 °C overnight and then purified using RP-HPLC with proteonavi column. The product XYZ-pS87/gS87 was lyophilized for next reaction. The segments after ligation were all characterized with analytical RP-HPLC and ESI-MS (Figure S3).

Radical catalyzed desulfurization was then performed to obtain α-syn protein with different modifications at serine 87. Briefly, about 2.6 mg of XYZ- pS87/gS87 was dissolve in 200μl of 6 M guanidine-HCl, 200 mM sodium phosphate buffer at pH 7.0 and mixed with 200 μl of 1 M TCEP and 40 μl of 2-methyl-2-propanethiol. 20 μl of 0.1 M 2-2’-azobis[2-(2-imidazolin-2-yl)propane] dihydrochloride (VA-044) was finally added to the mixture under argon. The reaction was shaken overnight at 37 °C. The pS87/gS87 α-syn protein was purified with proteonavi column, lyophilized, and characterized with analytical RP-HPLC and ESI-MS (Figure S4).

**Preparation of the unmodified WT, gS87 and pS87 α-syn fibrils and PFFs**

Recombinant unmodified WT, synthetic gS87 and pS87 α-syn in buffer containing 25 mM sodium phosphate (pH 7.4) were incubated at 37 °C, 900 rpm in ThermoMixer (Eppendorf) for 4 days with agitation in a concentration of 3.0 mg/ml, respectively. The fibril samples were further used for negative staining transmission electron microscopy, atomic force microscopy, cryo-EM sample preparation and rat primary neuron treatment. Unmodified WT, gS87 and pS87 α-syn PFFs were prepared by fibrils sonication at 20% power on ice for 25 times (1 s on, 1 s off).

**ThT kinetic assay**

50 μM unmodified WT, gS87 and pS87 α-syn monomers were incubated in 25 mM sodium phosphate (pH 7.4) buffer with or without unmodified WT α-syn PFFs in a black 384-well plate (Thermo Scientific). ThT was added to the reaction mixture at a final concentration of 30 μM. The fluorescence intensity was recorded using a Varioskan Flash Spectral Scanning Multimode Reader (Thermo Scientific), measuring at 440 nm (excitation) and 485 nm (emission) wavelengths with shaking at 900 rpm at 37 °C. Three replicates were performed for each sample. GraphPad Prism 9 was applied for graphing, with mean ± s.d..

**Negative staining transmission electron microscopy**

Five microliters of fibril solution were incubated on a 200-mesh glow-discharged copper grid (Zhongjingkeyi Technology Co., Ltd., Beijing) for 45 s. Then, the grid was washed with double-distilled water followed by 2% w/v uranyl acetate for another 45 s, and dried in air. The samples were imaged by a Tecnai T12 transmission electron microscope (FEI) operated in 120 kV.

**Atomic force microscopy**

Ten microliters of fibril solution were loaded on a clean mica surface for 5 min at room temperature and washed by double-distilled water into remove unbound fibrils. Next, images were captured by Nanoscope V Multimode 8 (Bruker) with SNL-10 probes (a constant of 0.35 N m^−1^) on ScanAsyst air mode in 1 Hz scan rate. The following data processes were carried out in the supplied software NanoScope Analysis (version 1.5, Bruker).

**Cryo-EM sample preparation and data collection**

The gS87 and pS87 α-syn fibril samples were applied into glow-discharged holey carbon copper grids (R2/1, 300 mesh, Quantifoil), and then plunge frozen in liquid ethane after blotting with filter paper by using Vitrobot Mark IV (Thermo Fisher).

Cryo-EM data of gS87 and pS87 was collected in a 300 kV Titan Krios G4 transmission electron microscope (Thermo Fisher) with a BioContinuum K3 direct detector (Gatan), using a GIF Quantum energy filter (Gatan) with a slit width of 20 e^-^V to remove inelastically scattered electrons. Movies with 40 frames per micrograph were recorded in ×105,000 magnification with a pixel size 0.83 Å pixel^-1^ at super-resolution mode. Cryo-EM data collection was performed by software EPU (Thermo Fisher) with 2 s exposure time and -1.0 to -2.0 μm defocus value in a total dose of 55 e^-^Å^2^.

**Image processing**

MotionCorr2^9^ for motion correction implement was carried out to correct beam-induced motion of movie frames with dose-weighting. Then, CTFFIND-4.1.8^10^ was used to estimate the contrast transfer function of motion-corrected images. Next, Fibrils were manually picked by using the manual picking method of RELION version 3.1^11^.

**Helical reconstruction**

Helical reconstruction was performed in RELION version 3.1^11^, including particle extraction, two-dimensional (2D) classification, 3D classification, 3D auto-refinement and post-processing.

For gS87 dataset, 21,328 manually fibrils from 2,134 micrographs were extracted into segments in a box-size of 360 pixels with an inter-box distance of 30.0 Å. Then, several iterations of 2D classification were applied to extracted segments with a decreasing in-plane angular sampling rate from 12° to 0.5° and the T = 2 regularization parameter. For double filament and single filament gS87 α-syn fibrils separated after 2D classification, a cylindrical map model generated by relion_helix_toolbox program was used as an initial 3D reference to perform the following 3D classification (K = 3). When the local search of symmetry for helical twist and rise was carried out after the separation of β-strands, the clearest class was selected for following 3D auto-refinement. Finally, the overall resolutions of double filament gS87 α-syn fibrils were reported at 3.1 Å, according to the gold-standard Fourier shell correlation (FSC) = 0.143 criteria.

For pS87 dataset, 27,806 fibrils from 2,423 micrographs were extracted into segments in a box-size of 360 pixels with an inter-box distance of 30.0 Å. Next, several iterations of 2D classification were applied to extracted segments with a decreasing in-plane angular sampling rate from 12° to 0.5° and the T = 2 regularization parameter. Similarly, for the twisted filament of pS87 α-syn fibril separated from 2D classification, a cylindrical map model generated by relion_helix_toolbox program was used as an initial 3D reference to perform the following 3D classification (K = 3). When the local search of symmetry for helical twist and rise was carried out after the separation of β-strands, the clearest class was selected for following 3D auto-refinement. Finally, the overall resolutions of twisted filament gS87 α-syn fibrils was reported at 2.6 Å, according to the gold-standard FSC = 0.143 criteria.

**Atomic model building and refinement**

Based on the density maps after post-processing program, the atomic models of double filament of gS87 α-syn fibril and twisted filament pS87 α-syn fibril were built de novo in COOT^12^, respectively. Then, three-layer models were generated in software Chimera and refined by real-space refinement program of PHENIX^13-14^. Additional details about two models were shown in Table S1.

**Primary neuronal cultures**

Embryonic day (E) 16 to E18 Sprague–Dawley rat (Shanghai SIPPR BK Laboratory Animals Ltd.) embryos were sacrificed for primary cortical neurons culture as previously described^15^. In brief, papain-digested cortical neurons were plated onto coverslips, coated with poly-L-lysine, in a 24-well plate at a density of 150,000 cells per well. After 7 days, neurons were treated with PBS, 100 nM WT or gS87 or pS87 α-syn PFFs, with three biological repetitions for each treatment. All animal experiments were performed according to the protocols approved by the Animal Care Committee of the Interdisciplinary Research Center on Biology and Chemistry (IRCBC), Chinese Academy of Sciences (CAS).

**Immunofluorescence staining and confocal imaging**

After treating for 14 days, neurons were collected for immunofluorescence imaging. Neurons plated on coverslips were fixed with 4% paraformaldehyde (PFA) and 4% sucrose in PBS and then permeabilized with 0.15% Triton X-100 diluted in PBS. Then coverslips were blocked with 3% goat serum (GS) diluted in PBS for 30 minutes at room temperature. Neurons were incubated with primary antibodies of phospho-α-synuclein (1:1,000, Abcam, 51253) and MAP2 (1:2,500, Abcam, 5392), diluted in 3% GS, at 4 °C overnight. After washing with PBST, 0.1 % Tween-20 diluted in PBS for three times, neurons were incubated with secondary antibodies of Alexa Fluor 488- and Alexa Fluor 568- (1:1,000, Invitrogen, 2420700, 2155282, respectively) for 1 h at room temperature. Without rinsing, DAPI stain was applied (1:10,000, Yeasen, 40728ES03) for 10 mins at room temperature. After rinsing, coverslips were mounted onto glass slides with mounting medium (ProLong Gold antifade reagent, Invitrogen, P36930). Confocal images were acquired by a laser scanning confocal microscope (SP8, Leica). A 63× water immersion objective was used for imaging. Images were batched analyzed by Image J. All images were projected in z direction with max intensity followed by background subtraction. The immunofluorescence results of the mean p-α-syn signal intensity were normalized to DAPI intensity of their own group.

**Cell viability assay**

Primary neurons were treated with different concentrations of unmodified WT, gS87 and pS87 α-syn PFFs or PBS after growing for 7 d. Three biological replicates were set up for each treatment. After treating for 14 d, the CCK-8 assay was performed following the manufacturer’s instructions. The absorbance at 450 nm wavelength was measured by a multimode plate reader (Ensight, PerkinElmer). Values were analyzed by GraphPad Prism 9.

**Supplementary Figures**

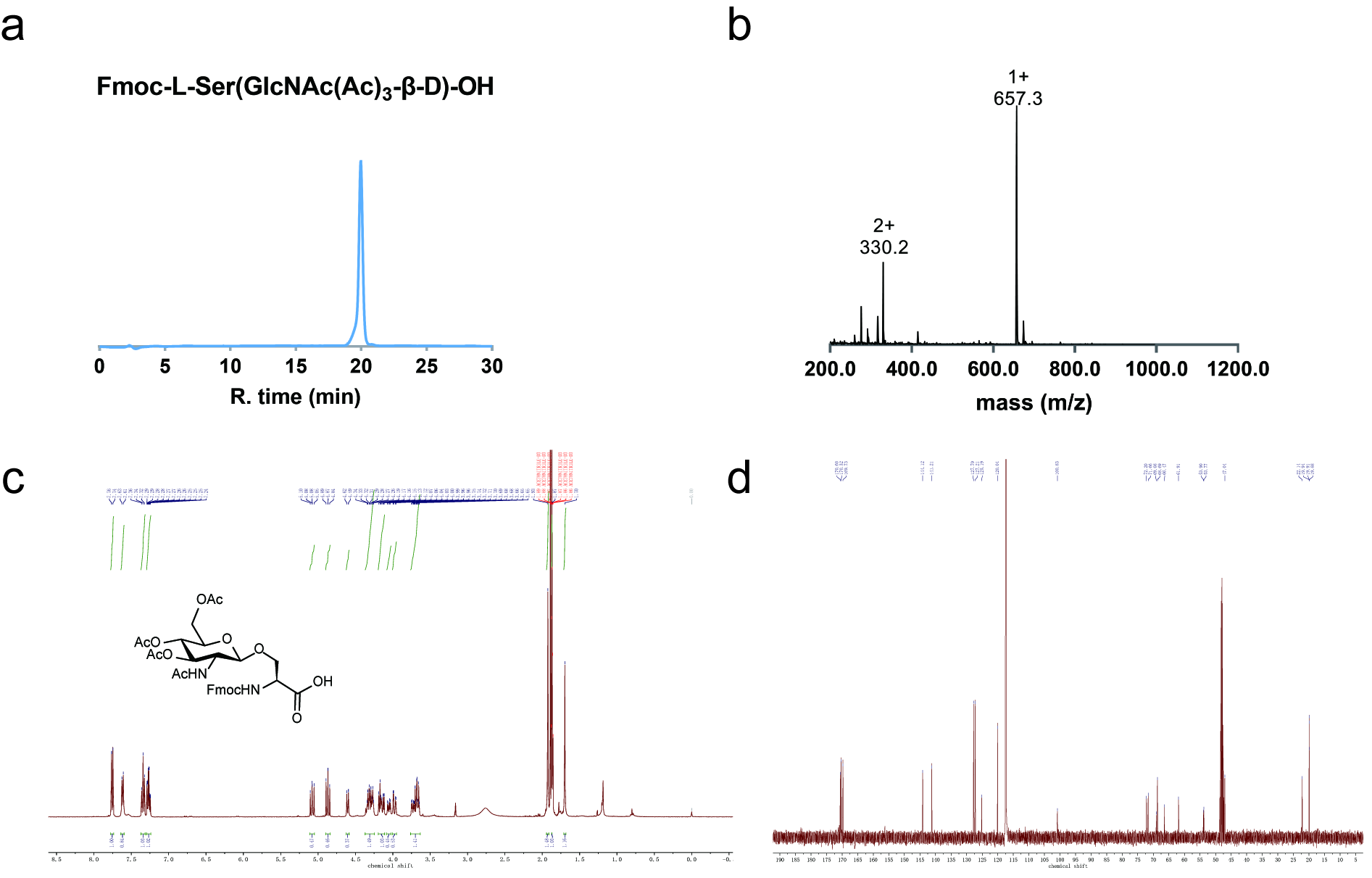

**Figure S1.** **Characterization of** **Fmoc-****L-Ser(GlcNAc(Ac)_3_-β-D)-OH.**

(a) Analytical HPLC result of Fmoc-L-Ser(GlcNAc(Ac)_3_-β-D)-OH. Retention time is about 19.95 min in a linear gradient of 50-90%B for 30 min (HPLC solvent A: water, 0.06% TFA; B: 80% CH_3_CN/water, 0.06% TFA). (b) ESI-MS characterization of Fmoc-L-Ser(GlcNAc(Ac)_3_-β-D)-OH. Calculated mass: 656.6 Da, observed mass: 656.3 Da. (c) ^1^H NMR characterization of Fmoc-L-Ser(GlcNAc(Ac)_3_-β-D)-OH (400 MHz, Acetonitrile-*d_3_*) δ 7.75 (d, J = 7.6 Hz, 2H), 7.62 (d, J = 7.5 Hz, 2H), 7.34 (t, J = 7.5 Hz, 2H), 7.27 (tdd, J = 7.4, 3.6, 1.2 Hz, 2H), 5.08 (dd, J = 10.6, 9.4 Hz, 1H), 4.95 – 4.80 (m, 1H), 4.60 (d, J = 8.5 Hz, 1H), 4.37 – 4.25 (m, 3H), 4.20 – 4.11 (m, 2H), 4.05 (dd, J = 10.5, 4.6 Hz, 1H), 3.98 (dd, J = 12.3, 2.5 Hz, 1H), 3.76 – 3.63 (m, 3H), 1.93 (s, 3H), 1.87 (s, 2H), 1.70 (s, 2H). (d) ^13^C NMR characterization of Fmoc-L-Ser(GlcNAc(Ac)_3_-β-D)-OH (400 MHz, Acetonitrile-*d_3_*) δ 170.60, 170.32, 169.73, 144.12, 141.21, 127.79, 127.21, 125.19, 120.04, 100.83, 72.20, 71.66, 69.08, 68.69, 66.47, 61.91, 53.90, 53.77, 47.04, 22.14, 19.94, 19.91, 19.88.

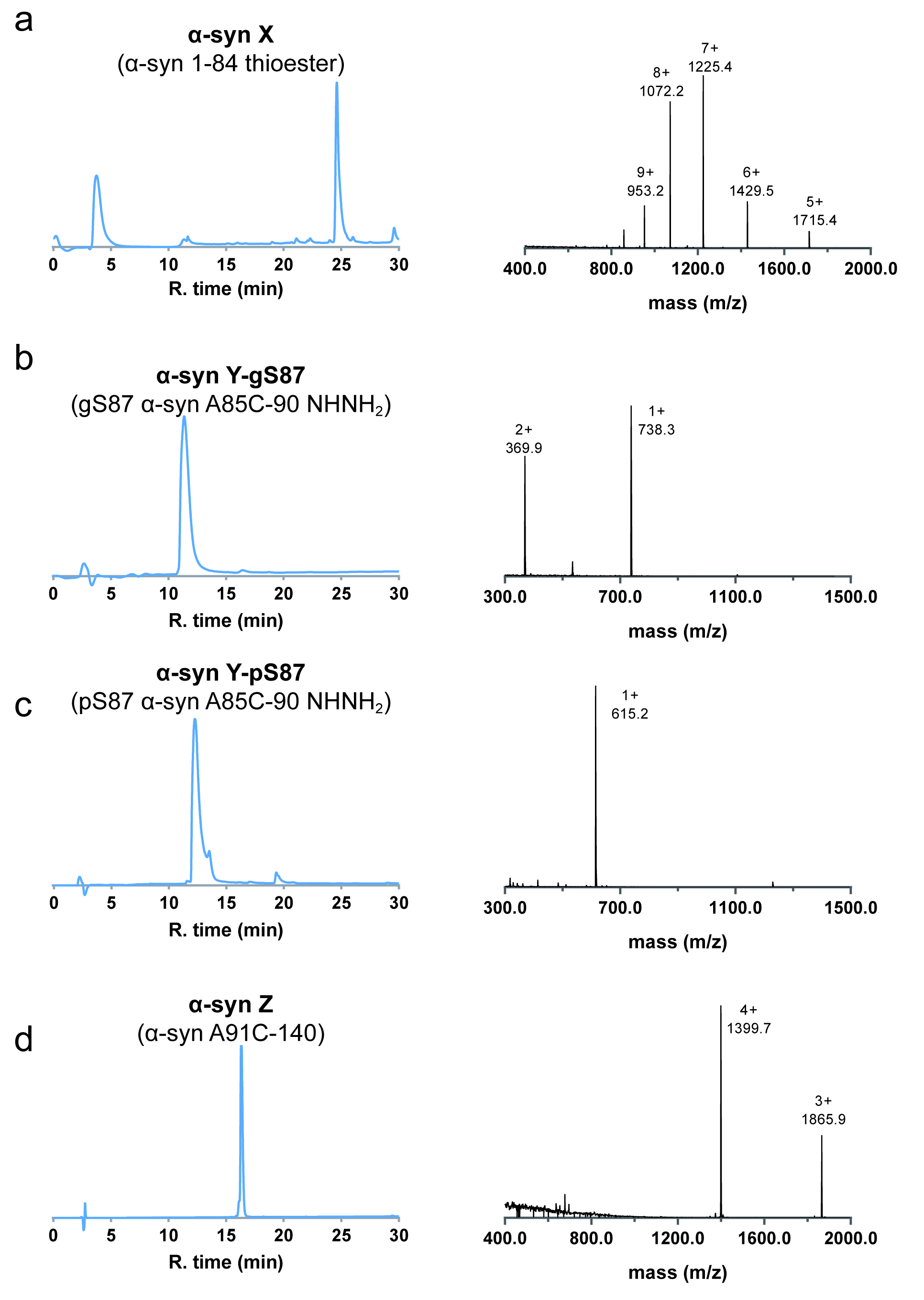

**Figure S2. Characterization of the synthetic segments: X, Y, and Z.**

(a) Characterization of α-syn X (MesNa thioester). Left: analytical HPLC result of α-syn X in Buffer. Retention time is about 24.62 min in a linear gradient of 30-70%B for 30 min; Right: ESI-MS characterization of α-syn X (MesNa thioester). Calculated mass: 8569.7 Da, observed mass: 8570.2 Da. (b) Characterization of α-syn Y-gS87. Left: analytical HPLC result of α-syn Y-gS87. Retention time is about 11.36 min in a linear gradient of 5-25%B for 30 min; Right: ESI-MS characterization of α-syn Y-gS87. Calculated mass: 737.3 Da observed mass: 737.5 Da. (c) Characterization of α-syn Y-pS87. Left: analytical HPLC result of α-syn Y-gS87. Retention time is about 12.25 min in a linear gradient of 5-25%B for 30 min; Right: ESI-MS characterization of α-syn Y-pS87. Calculated mass: 614.2 Da, observed mass: 614.2 Da. (d) Characterization of α-syn Z. Left: analytical HPLC result of α-syn Z. Retention time is about 16.31 min in a linear gradient of 30-70%B for 30 min; Right: ESI-MS characterization of α-syn Z. Calculated mass: 5594.0 Da, observed mass: 5594.7 Da.

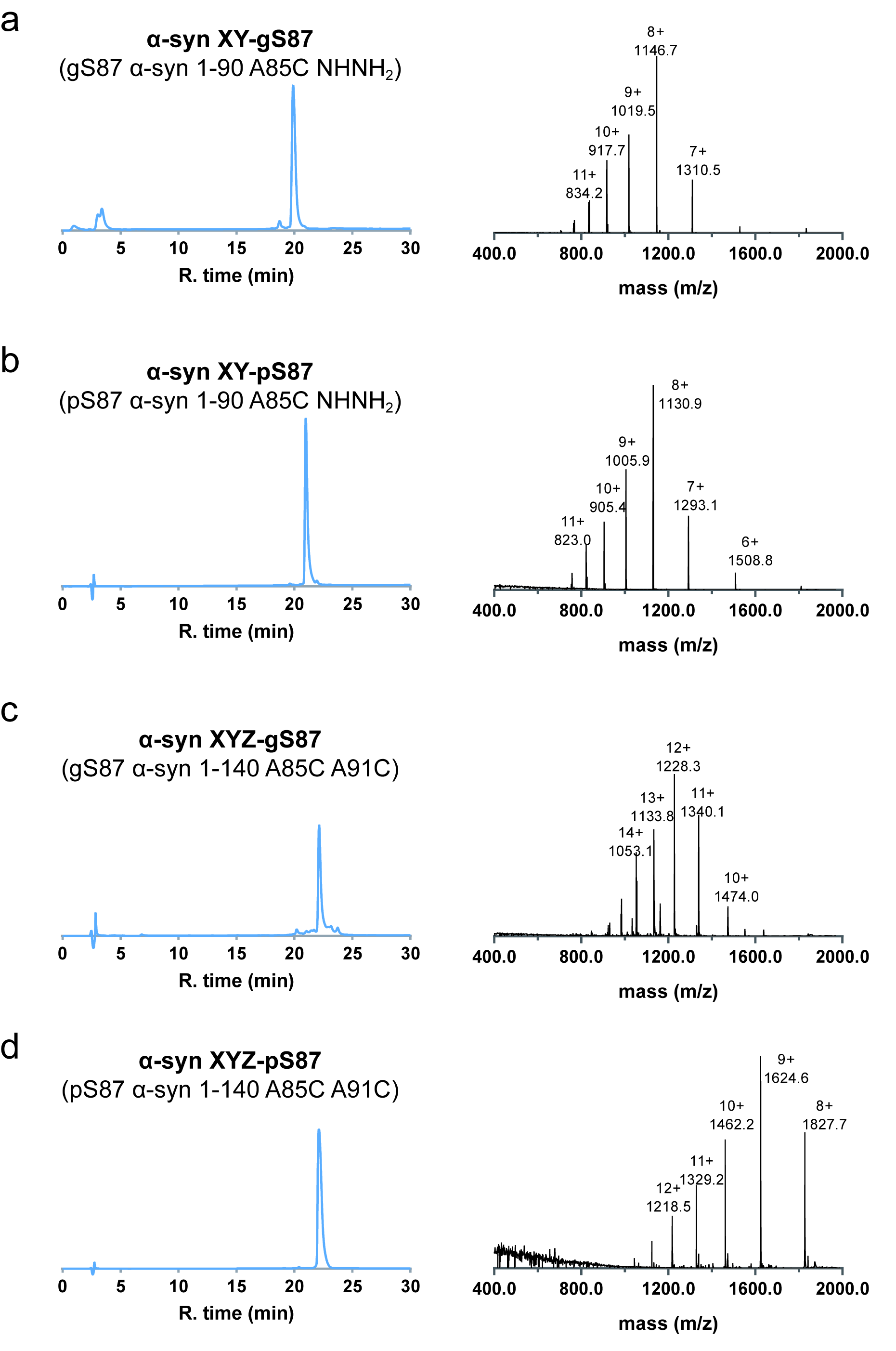

**Figure S3. Characterization of the synthetic segments: XY and XYZ.**

(a) Characterization of α-syn XY-gS87. Left: analytical HPLC result of α-syn XY-gS87. Retention time is about 19.88 min in a linear gradient of 30-70%B for 30 min; Right: ESI-MS characterization of α-syn XY-gS87. Calculated mass: 9165.4 Da observed mass: 9165.8 Da. (b) Characterization of α-syn XY-pS87. Left: analytical HPLC result of α-syn XY-pS87. Retention time is about 20.98 min in a linear gradient of 30-70%B for 30 min; Right: ESI-MS characterization of α-syn XY-pS87. Calculated mass: 9042.3 Da, observed mass: 9043.7 Da. (c) Characterization of α-syn XYZ-gS87. Left: analytical HPLC result of α-syn XYZ-gS87. Retention time is about 22.13 min in a linear gradient of 30-70%B for 30 min; Right: ESI-MS characterization of α-syn XYZ-gS87. Calculated mass: 14727.3 Da, observed mass: 14729.2 Da. (d) Characterization of α-syn XYZ-pS87. Left: analytical HPLC result of α-syn XYZ-pS87. Retention time is about 22.11 min in a linear gradient of 30-70%B for 30 min; Right: ESI-MS characterization of α-syn XYZ-pS87. Calculated mass: 14604.3 Da, observed mass: 14612.1 Da.

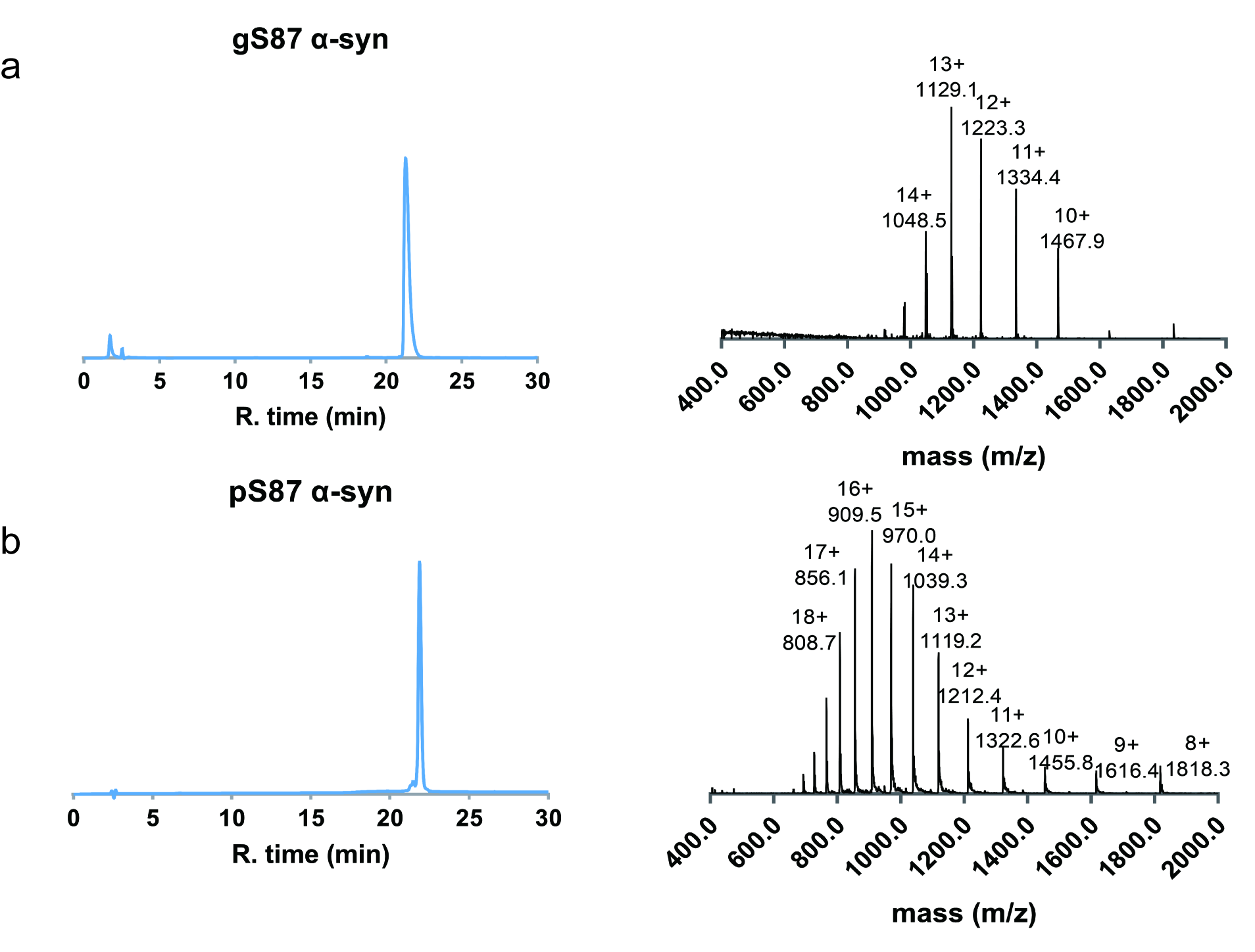

**Figure S4. Characterization of the synthetic pS87 α-syn and gS87 α-syn.**

(a) Characterization of gS87 α-syn. Left: analytical HPLC result of gS87 α-syn. Retention time is about 21.28 min in a linear gradient of 30-70%B for 30 min; Right: ESI-MS characterization of gS87 α-syn. Calculated mass: 14663.3 Da, observed mass: 14666.1 Da. (b) Characterization of pS87 α-syn. Left: analytical HPLC result of pS87 α-syn. Retention time is about 21.46 min in a linear gradient of 30-70%B for 30 min; Right: ESI-MS characterization of pS87 α-syn. Calculated mass: 14540.1 Da observed mass: 14536.4 Da.

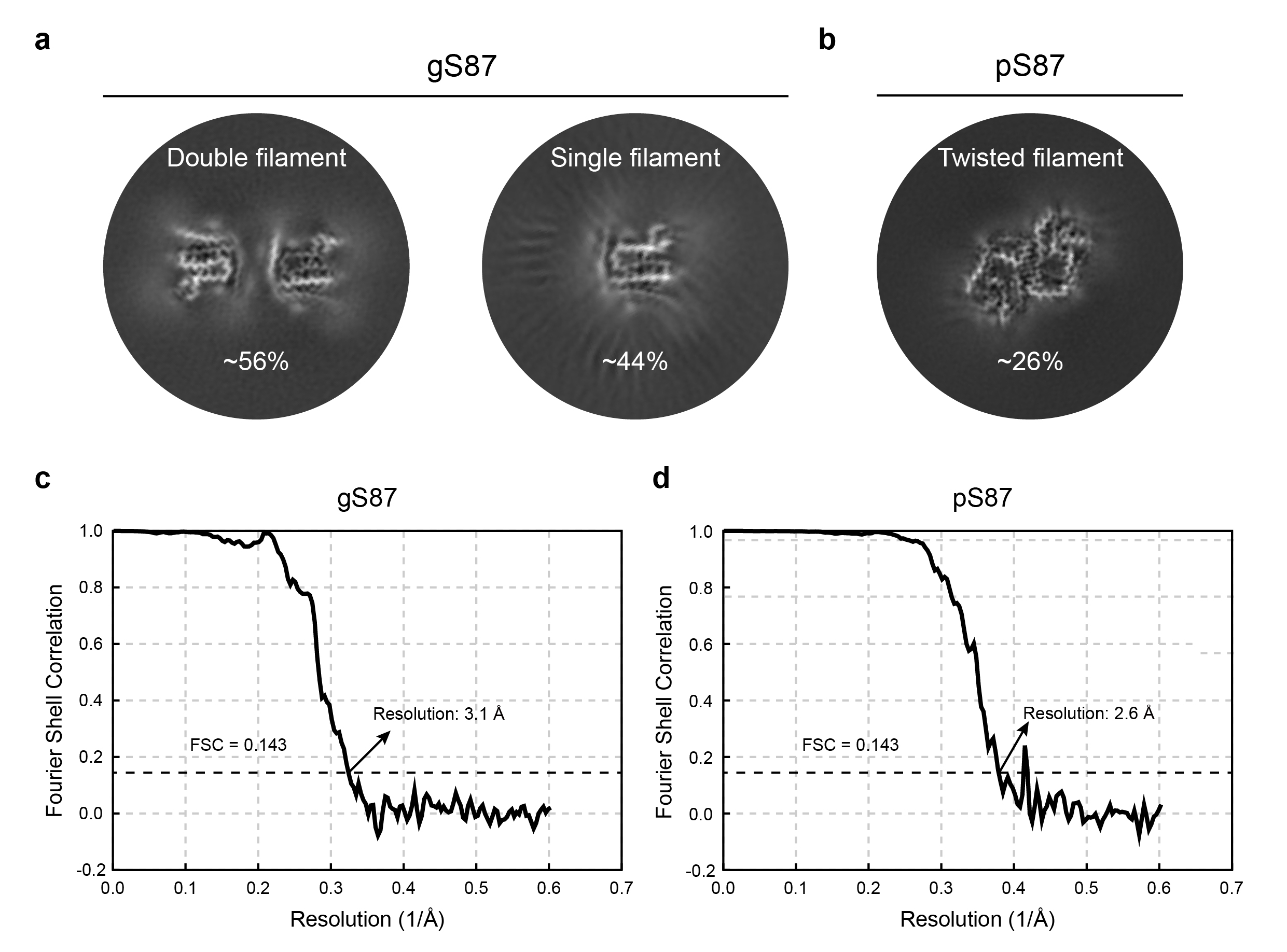

**Figure S5. Cryo-EM structure determination of the gS87 and pS87 α-syn fibrils.**

(a) 3D classification results of the double filament polymorph (~56%) and the single filament polymorph (~44%) of the gS87 α-syn fibril. (b) 3D classification result of the twisted filament polymorph (~26%) of the pS87 α-syn fibril. (c-d) Gold-standard Fourier shell correlation (FSC) curves of the density maps of the gS87 (c) and pS87 (d) α-syn fibrils.

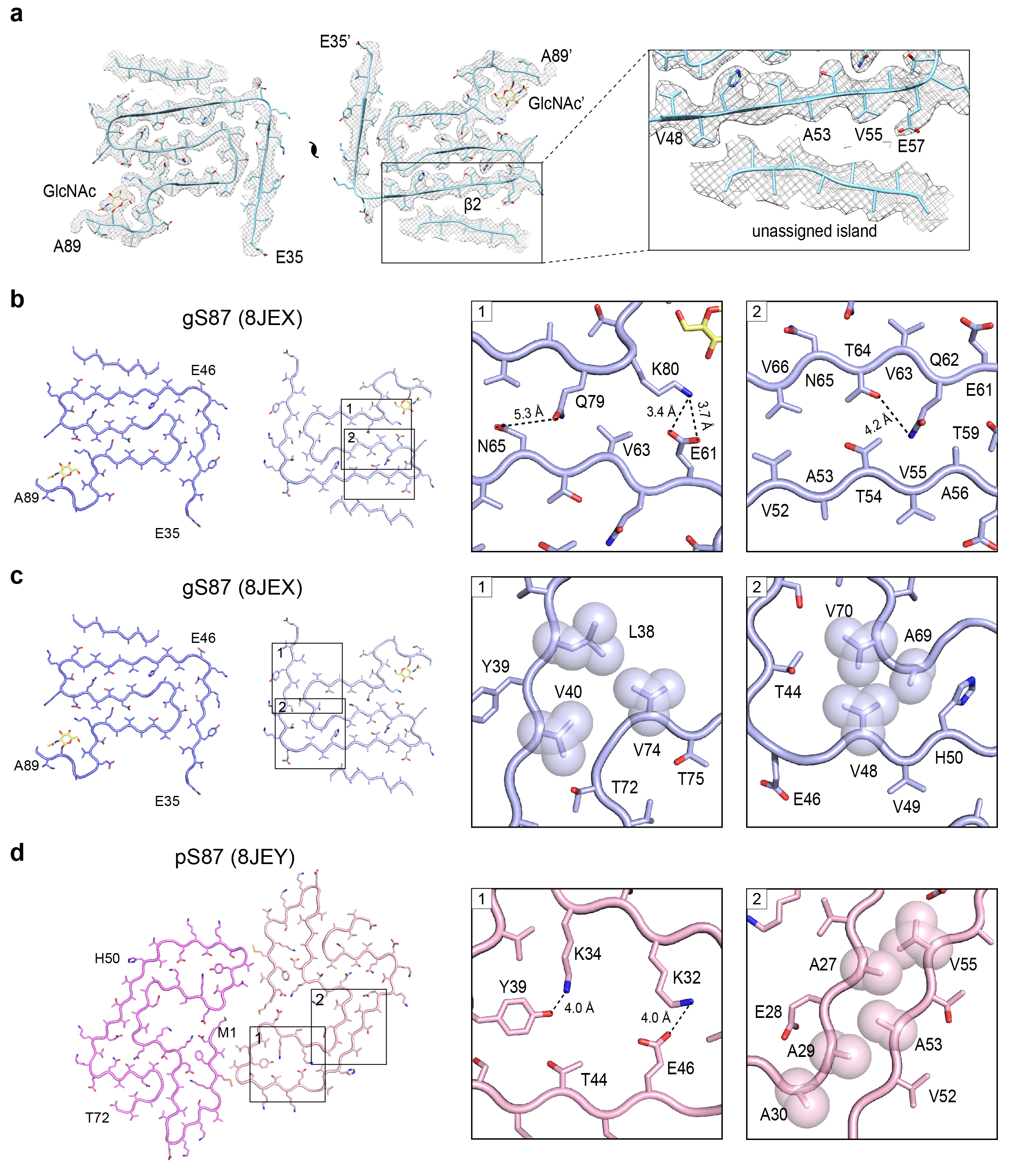

**Figure S6. Structural analysis of the gS87 and pS87 α-syn fibrils.**

(a) In the electron density map of the gS87 fibril, the unassigned island was observed on the outer surface of the fibril core, which was adjacent to β2. (b) Zoom-in views of hydrophilic zipper-like interactions in gS87 fibril structure to the stabilization of the U-shaped structure. (c) Zoom-in views of the hydrophobic interactions in gS87 fibril model. Residues involved in the interactions are indicated in spheres. (d) The structure of the pS87 fibril, with the salt bridge between K32 and E46, the hydrogen bond between K34 and Y39, and the steric zipper-like hydrophobic interaction shown in the zoom-in views.

**Table S1.** Cryo-EM data collection, modeling and refinement statistics.

| **Data collection and processing** | **gS87 α-syn**  (EMD: 36202)  (PDB: 8JEX) | **pS87 α-syn**  (EMD: 36203)  (PDB: 8JEY) |
| --- | --- | --- |
| **Data Collection**  Magnification (×) | 105,000 | 105,000 |
| Pixel size (Å) | 0.83 | 0.83 |
| Defocus Range (μm) | -1.0 to -2.0 | -1.0 to -2.0 |
| Voltage (kV) | 300 | 300 |
| Camera | BioContinuum K3 | BioContinuum K3 |
| Microscope | Krios G4 | Krios G4 |
| Exposure time (s/frame) | 0.05 | 0.05 |
| Number of frames | 40 | 40 |
| Total dose (e^-^/Å^2^) | 55 | 55 |
| **Reconstruction** |  |  |
| Micrographs | 2,134 | 2,423 |
| Manually picked fibrils | 21,328 | 27,806 |
| Box size (pixel) | 360 | 360 |
| Inter-box distance (Å) | 30 | 30 |
| Initial particle images (no.) | 465,930 | 647,678 |
| Final particle images (no.) | 24,910 | 61,047 |
| Resolution (Å) | 3.1 | 2.6 |
| Map sharpening B-factor (Å^2^) | -96.5004 | -86.4792 |
| Helical rise (Å) | -179.72 | -179.72 |
| Helical twist (°) | 2.41 | 2.41 |
| **Atomic model** |  |  |
| Non-hydrogen atoms | 2,616 | 3,060 |
| Protein residues | 384 | 432 |
| Ligands | 6 | 0 |
| r.m.s.d. Bond lengths | 0.003 | 0.003 |
| r.m.s.d. Bond angles | 0.647 | 0.650 |
| All-atom clash score | 8.39 | 10.60 |
| Rotamer outliers | 0 % | 0 % |
| Ramachandran Outliers | 0 % | 0 % |
| Ramachandran Allowed | 3.33 % | 5.71 % |
| Ramachandran Favored | 96.67 % | 94.29 % |
